# Genome misassembly detection using Stash: A data structure based on stochastic tile hashing

**DOI:** 10.1101/2025.09.19.677272

**Authors:** Armaghan Sarvar, Lauren Coombe, Inanc Birol

## Abstract

**Motivation:** Analyzing large data from high-throughput sequencing technologies presents significant challenges in terms of memory and computational requirements. It is crucial to develop efficient data structures and computational methods that handle sequencing information. These challenges impact bioinformatics studies, including *de novo* genome assembly which serves as the foundation of genomics. Issues like read errors or limitations of heuristic decisions in assembly algorithms lead to genome misassemblies and inaccurate genomic representations, compromising the quality of downstream analyses. Hence, *de novo* assemblies can benefit from misassembly detection and correction, to produce a more optimal assembly.

**Results:** We present Stash, a novel hash-based data structure designed for storing and querying large sequencing data. For an input sequence, Stash uses sliding windows of spaced seed patterns to extract and hash *k*-mers. The hash values combined with the sequence ID determine the value stored in Stash. A filled Stash can be used to query whether two genomic regions are covered by the same set of reads. This can be used in genome misassembly detection. We demonstrate the effectiveness of Stash in detecting misassemblies in human genome assemblies generated by Flye and Shasta, using Pacbio HiFi reads from the human cell line NA24385. We observed that scaffolding Stash-cut assemblies reduces 7.6% and 3.4% of misassemblies in the Flye and Shasta assemblies, respectively. This is accomplished in 310 minutes utilizing 8 GB of memory. Stash is comparable to alternative long read misassembly correction methods and can result in superior assemblies compared to the baseline.

**Availability:** Stash is implemented in C++ as a static library and is available here: https://github.com/bcgsc/Stash.

## 1. Introduction

The rapid advancements in high-throughput sequencing technologies have opened new avenues to be explored using bioinformatics, such as the discovery of complex genomic landscapes or the identification of genetic variations. With associated massive improvements in sequence throughput, storing and analyzing the extracted Gbp-scale genomic information has become a major logistical and computational challenge. Hence, novel data structures and algorithms borrowed from the broader Computer Sciences have been developed to fuel rapid development in addressing these challenges (Brown & Morgenstern, 2014).

An example is the Bloom filter (Bloom, 1970), which is a probabilistic and memory-efficient data structure designed for fast membership queries. Bloom filters can represent large quantities of data succinctly by hashing values and using them as indices in a bit array. In recent years, Bloom filters have been improved to extend their practical utility in scientific research. An example is the multi-index Bloom filter (miBF) data structure (Chu, et al., 2020), which stores a vector of identifiers along with the membership bits, allowing for applications like taxonomy assignment and sequencing read binning without the need for sequence alignment, a costly bioinformatics task in terms of run time and memory. Other advanced data structures used in this field include the quotient filter (Bender, et al., 2011), which utilizes hashing and division operations to store and query data, and the counting quotient filter (Pandey, et al., 2017) that also allows efficient counting of the frequency of each element. Cuckoo filters (Fan, et al., 2014) are another improvement over Bloom filters that resolve hash collisions by evicting existing elements from their original locations. There is also a probabilistic indexing data structure called the quasi dictionary (Marchet, et al., 2020), designed based on a Minimal Perfect Hash Function, to provide a way to associate any kind of data to a given input key.

Many bioinformatics algorithms such as structural variation detection (Ryan P. Abo, et al., 2015) (Wang, et al., 2021) or protein function and structure prediction (Hippe, et al., 2020) (Qin, et al., 2012) are based on identifying short, fixed-length substrings of reads, which are called *k*-mers. The *k*-mer based methods work by indexing sequence substrings with length *k*, usually in a hash-based data structure. In the querying phase, a given sequence is also segmented into *k*-mers and searched against the index to find shared subsequences. Moreover, hash-based data structures are frequently used to count consecutive *k*-mers for tasks such as similarity searching in large-scale analysis (Marchet, et al., 2021). Accordingly, using high performance hashing algorithms would have a substantial effect on the deployment of these data structures (Mian, et al., 2023). An example is ntHash2 (Kazemi, et al., 2022), which is a recursive hashing algorithm based on ntHash (Mohamadi, et al., 2016) tuned for processing DNA/RNA sequences, taking advantage of the *K ―*1 overlap between consecutive *k*-mers.

However, *k*-mer based representations are not immune to sequencing errors. Hence, some tools, such as the miBF, use spaced seed patterns represented by sequences of zeroes and ones, with ones indicating care and zeroes indicating don’t care positions. Spaced seeds can thus tolerate mismatches, allowing certain base pairs defined by the pattern to mismatch without penalty.

Among the challenges in the domain of high-throughput sequencing analysis and big-data-driven life sciences, *de novo* genome assembly is one of the fundamental unsolved problems to address. Despite recent improvements in long read sequencing technologies, assembling large genomes without a reference genome remains a challenging task for assembly algorithms. *De novo* genome assembly has led to the generation of high-quality reference genomes (Dida & Yi, 2021), and it has many applications such as in phylogenetic inference (Fitz-Gibbon, et al., 2017), gene annotation (Warren, et al., 2015), and cancer genomics (Park & Kim, 2016). However, because of genome repeats and complexities, errors in sequencing reads, and assembly pipeline artifacts, contig misassemblies are common in most genome sequencing projects. Hence, an important problem in *de novo* genome assembly is the detection and correction of draft genome misassemblies, leading to improved overall assembly quality.

To discover misassemblies in *de novo* assemblies, some identification methods have been proposed, and they are mostly based on aligning the sequencing reads back onto the assembly to find measurements for assembly quality. Tools such as REAPR (Hunt, et al., 2013) or Pilon (Walker, et al., 2014) use short reads to carry out misassembly detection using read to contig alignments. REAPR utilizes a scoring system that assesses both local accuracy and the presence of larger-scale errors within assemblies. In this approach, error-free bases correspond to regions of that are likely to be accurate. Conversely, bases failing to meet these criteria receive scores ranging from zero to one, determined by the deviation from acceptable thresholds in metrics such as read depth, characteristics of paired mapping, and the presence of soft clipping. Pilon, considers assembly improvements (polishing and error correction) and variant detection as a unified process, starting with an input assembly and employing read alignments to pinpoint discrepancies. In regions with suboptimal alignments, Pilon conducts local reassembly, filling gaps, and capturing large-scale insertions and deletions. This approach encompasses the identification and rectification of base errors and the detection of potential local misassemblies.

Linked reads, a sequencing technology developed by 10x Genomics (Pleasanton, CA, US) for their Chromium platform can also be used to correct misassemblies, such as in Tigmint (Jackman, et al., 2018). Tigmint identifies drops in molecule coverages inferred from the alignment of linked reads to the assembly. Since linked reads can provide versatile long-range information, they can detect genome misassemblies better than short reads (Peona, et al., 2021).

However, long reads from the third-generation sequencing technologies provide the most comprehensive view of genomic regions. Tools such as Tigmint-long (Coombe, et al., 2021), ReMILO (Bao, et al., 2018), Inspector (Chen, et al., 2021), and GAEP (Zhang, et al., 2023) have been proposed as long-read misassembly detection tools. Tigmint-long is based on the Tigmint tool explained above and has been adapted to work with long reads. ReMILO is a reference-assisted misassembly detection algorithm that leverages both short reads and PacBio SMRT long reads, utilizing their complementary strengths rather than being exclusive to long-read information. Similar to Tigmint-long, in the Inspector and GAEP tools, sequencing reads are aligned to the assembly. Inspector detects and corrects errors in the assembly by first inferring assembly and read-to-contig alignment statistics such as the read mapping rate and average alignment depth, and then using several rounds of alignment and local assembly to correct the detected misassembly regions. GAEP does not output a corrected assembly, and for reporting detected misassemblies, it leverages the fact that long-read alignments can break at the corresponding misassembly positions. It is worth noting that these tools rely on long-read alignments, which are particularly computationally intensive to generate, especially when working with large genomes.

Here, we introduce stochastic tile hashing (Stashing) to describe large collections of sequences through a lossy and scalable representation. Our proposed novel underlying data structure, Stash, uses a fixed allocated memory, where the memory loci are indicated by some hashed values of subsequences, and the stored value is partly determined by the index of the sequence. The Stash data structure efficiently facilitates saving and retrieving sequence mapping information.

In contrast to membership data structures such as the Bloom filter which let us store and query information regarding the presence or absence of a set of observations, the Stash data structure also provides us with the utility of implicitly storing a property that corresponds to the input observations, such as sequencing reads. When querying the data structure, we can compare the stored property for two observations of interest. For example, Stash can determine if two sequences originate from similar genomic regions without the need for computationally intensive sequence alignment. We illustrate the utility of Stash in detecting and correcting genome misassemblies, focusing on the use of PacBio HiFi sequencing reads. Due to its effectiveness in storing large amounts of regional information, we expect the Stash data structure to have broad applications in other areas of genomics research as well, such as scaffolding and polishing.

## 2. Methods

### 2.1 Filling the Stash Data Structure

The underlying structure of Stash is a bit array represented in two dimensions where the rows have *t*_1_ tiles, each with *t*_2_ bits. Given a set of sequences, the Stash data structure is populated as shown in Figure 1A. First, for each input sequence, the associated sequence ID is hashed twice using two different hash functions, and each value is segmented into multiple tiles. Second, each input sequence is repeatedly hashed with sliding windows of symmetric spaced seed patterns. We use a set of *h* spaced seed patterns each having a weight of *w* and length of *k*. Next, the hashed values extracted after applying ntHash2 determine which rows of the Stash memory block to update. The sequence hashing function used in filling and querying Stash needs to be canonical when dealing with genomics information, meaning it produces identical values for a sequence and its reverse-complement, reducing the necessary storage space for sequences by half. With *h* spaced seed patterns, each sliding window position updates *h* rows of Stash, and we refer to each of these *h* accessed rows as a Stash frame. Using more than one spaced seed pattern will reduce the ntHash collision possibilities, reducing the probability of different *k*-mers getting mapped to the same Stash frame.

**Fig. 1.**
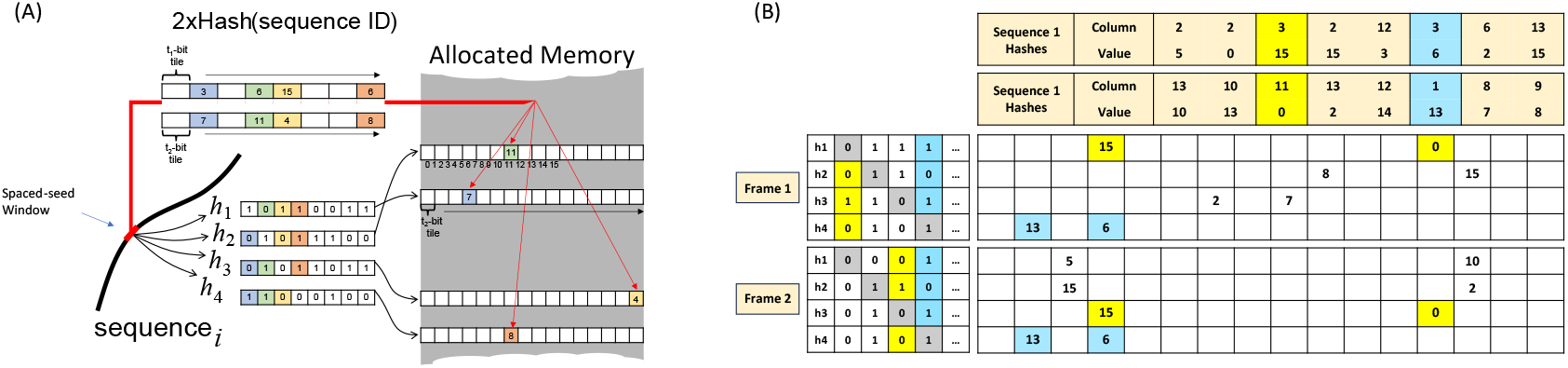
The Stash data structure. (A) Algorithm, data structure and sequencing data population process with four spaced seed patterns (h1-h4). (B) The results of sample frames after filling Stash with two overlapping sequencing reads. The colors show equal hash outputs resulting in the same columns and tile values extracted from the sequence ID for the overlapping regions.

For each spaced seed pattern, to determine the destination tile and its content to be stored in the selected Stash frame row, the hashes corresponding to the other spaced seed patterns are combined with the two sequence ID hash values aforementioned. More specifically, for the *i*th spaced seed pattern and its corresponding hash value, the *i*th most significant bits of the other hash values are concatenated to create an index addressing the tiles in the two sequence ID hashes. One addressed tile specifies the column, while the other indicates the value to be stored in a Stash tile. As an example of how Stash is filled, we populate the data structure with two sequencing reads and show how it leads to repeated patterns when the two spaced seed sliding windows fall in an underlying region where the sequences overlap (Figure 1B).

### 2.2 Querying the Stash Data Structure

To query the Stash data structure and answer the question of whether two genomic regions are potentially covered by the same set of sequencing reads, we define a score metric called the “number of matches”. This metric captures the similarity between any two populated Stash frames, or two windows composed of frames.

#### 2.2.1 The Number of Matches for Two Frames

Based on the Stash filling process, for any two spaced seed pattern outputs coming from the same input sequence, we expect the two corresponding Stash frames to contain a number of equal tile values when examining the two frame columns extracted with the same index. More specifically, it is likely to observe equal pairs of column and tile values when comparing their corresponding Stash frames, as they access the same sequence ID hash vectors when being inserted into Stash.

Hence, the number of matches similarity metric for a pair of frames is calculated by counting the tiles with equal values in each Stash frame column. Specifically, the number of matches between two frames is incremented if a tile in one frame has the same value as at least another tile in the corresponding column of the second frame. This ensure commutative property when counting the number of matches between two Stash frames.

#### 2.2.2 The Number of Matches for Windows of Frames

To have a more general view of a genomic region, we define windows of Stash frames. A window consists of a specific *number of frames* each starting at a distance called *stride* of the starting point of the previous frame. Any two sliding windows of frames have an adjustable distance of *delta*. Figure 2 shows a diagram of two windows placed over a sequence.

**Fig. 2.**
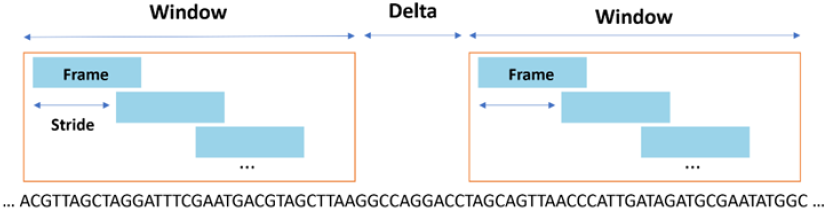
A pair of windows of Stash frames.

For two windows of frames, we calculate the aforementioned number of matches score for every pair of frames residing in the windows and take the maximum score among all, which demonstrates how likely it is that the underlying sequences of the two windows originate from the same genomic region. Supplementary Figure 1 demonstrates the effectiveness of this metric in an arbitrary case.

### 2.3 Stash Properties

#### 2.3.1 Saturation

The Stash data structure can get saturated if the same *k*-mers are repeatedly added into it. This is particularly relevant when the sequence coverage depth is high in a given read set. We define saturation as the probability of a row being filled completely by the resulting hashed values of spaced seed pattern outputs. Assuming Stash has *C* columns, the probability of inserting into a row tile would be 1/*c*. If we define the number of insertions of the same spaced seed as *i*, the Stash row occupancy would be based on the following formula (Equation 1).

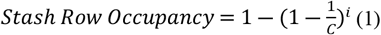

#### 2.3.2 Time Complexity

For an input sequencing read set of average length *l*_*avg*_ and *n*_*r*_ reads, assuming ntHash2 is used, the time complexity of filling the data structure would be as follows.

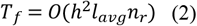

where *h* is the number of spaced seed patterns applied to the sequence before using ntHash, i.e., the number of frame rows. Here, we can see that the time complexity of filling Stash is linear with respect to the size of the input data.

On the other hand, considering two input *k*-mers and their corresponding extracted frames, the time complexity of comparing the two frames by hashing the sequences and counting the number of matches would be

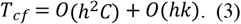

Additionally, when comparing two windows, i.e., sets of frames, as all pairs of frames will be examined at a time, by defining the *number of frames* in each window as *f*, and assuming that *h* × *c* is larger than *K*, the time complexity would be

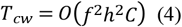

### 2.4 Genome Misassembly Detection and Correction

After populating the Stash data structure with a set of sequencing reads based on the steps explained in Section 2.1, having the corresponding input genome assembly of the reads, we extract the matches signal of each assembly contig by sliding a pair of Stash windows across the contig and computing the number of matches metric for each position. Next, we apply the max pooling operation over the extracted signal to reduce the noise and disregard the unexpected low number of matches, which arise due to genome repeats and complexities or assembly method artifacts. More specifically, a window is defined with the size of the pooling region, which specifies the number of consecutive bases pairs which will be compared together. It is then slid over the signal, extracting the maximum value within each region, and this process is repeated until the entire signal is covered.

Next, we compare each value of the generated signal with a threshold specified based on the expected number of matches between two unrelated windows of frames. If the value is less than the defined threshold, Stash reports a misassembly position since the two inspected windows of frames are likely not covered by the same set of sequencing reads and the underlying sequences do not originate from neighboring genomic regions. As a result, Stash will cut the assembly at the detected positions with low number of matches values using a module called *StashCut*. Breaking misassemblies allows for the possibility of the resulted sequences being correctly joined during scaffolding.

### 2.5 Experiments

Our experiments show how Stash is able to probabilistically infer whether two genomic regions originate from the same set of sequencing reads. For demonstration, we consider a Stash with 2^30^ rows and 16 four-bit tiles and fill it with ~30-fold coverage Pacbio HiFi long sequencing reads from the human cell line NA24385 (Wenger, et al., 2019) (Supplementary Table 1). The Stash dimensions were chosen in order for the Stash frames to be saturated based on the coverage of the input readset (refer to Equation 1), while also limiting the total memory usage of Stash to 8 GB, a value small enough to fit on many consumer-level computers. We use four symmetrical spaced seeds of length 26 and weight 18 to hash each sequence *k*-mer. The number of spaced seeds used balances ensuring robustness to mismatches without dedicating too much memory for any single Stash frame.

For the first experiment, we define a set of window distances (*delta*). Then, for each distance, we randomly choose 10,000 positions on the read set, a quantity that we empirically found large enough to ensure a robust representation of the distribution. This results in a match distribution for each *delta*. We also prepare a 10,000-point distribution for completely unrelated windows which can be conceptualized as a *delta* of *infinity*. This was simulated by randomly placing the two windows over reads of different chromosomes.

To showcase the performance of Stash in misassembly detection, we analyze human genomes assembled by Flye (Kolmogorov, et al., 2019) and Shasta (Shafin, et al., 2020) using the same read set with which we filled the Stash. Both assemblies are also scaffolded using ntLink (Coombe, et al., 2023), a recent long-read genome scaffolder, to give us a total of four baseline assemblies (refer to Supplementary Table 2 for details of each assembly). On both Flye and Shasta assemblies, we explored 5^5^ = 3,125 parameter configurations of *StashCut*, representing a coarse grid search over five parameters each with five possible values, as described in Supplementary Method 1. We then perform ntLink scaffolding on each run and use QUAST (Gurevich, et al., 2013) to evaluate the final assemblies, utilizing the reference genome GRCh38 to generate statistics such as contiguity metrics and the number of misassemblies.

Finally, a default parameter set for *StashCut* is chosen as discussed in the Results section and used to compare the performance of Stash-based misassembly correction with Tigmint-long and Inspector.

All benchmarking tests have been performed on a server-class system with 128 Intel(R) Xeon(R) CPU E7-8867 v3 @ 2.50 GHz with 2.4 TB RAM.

## 3. Results

As Δ increases, the match distributions between window pairs converge toward the distribution for unrelated windows (Δ = ∞), indicating the windows are more likely from non-adjacent genomic regions; operationally, we mark such cases using the 90% prediction-interval threshold derived from the unrelated distribution (Figure 3). This experiment demonstrates the capability of the Stash data structure to distinguish between related and unrelated windows within a queried sequence. As detailed in Supplementary Table 3, for delta values less than 2^11^ = 2,048, between 80% to 90% of all matches are already higher than the 90% prediction interval threshold (the drawn red line). Supplementary Table 4 shows for delta values less than 512, between 70% to 80% of all matches are higher than the 99% prediction interval threshold.

**Fig. 3.**
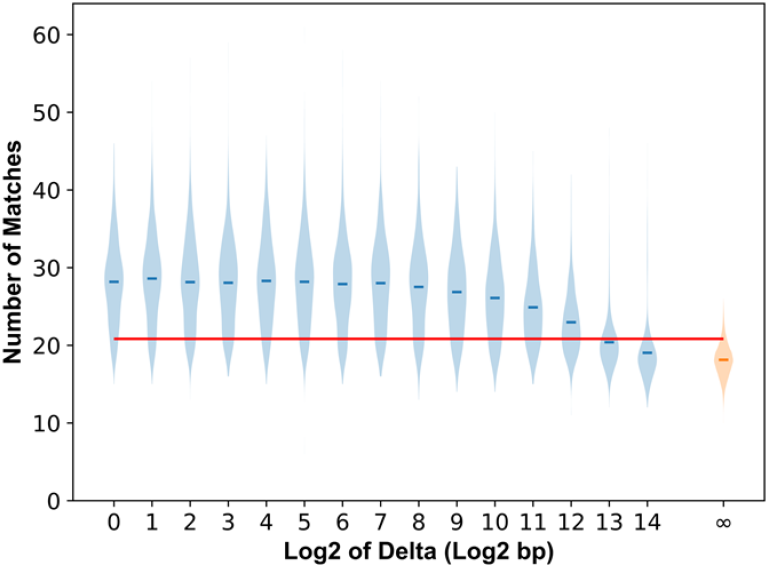
The distribution of the number of matches between two windows of frames as we increase the distance between them. Distance of infinity represents windows from sequencing reads of different chromosomes.

The *StashCut* misassembly detection is controlled by five parameters, namely *number of frames, stride, delta, threshold*, and *kernel radius*.

Figure 4 shows that many explored configurations of *StashCut* from the parameter space of Table 1 will either overcut the Flye assembly (the bottom left region) and therefore have low misassemblies and low contiguity represented by NGA50 or undercut and leave both metrics high (the top right region). There are, however, configurations that are superior to the others, considerably reducing extensive misassemblies without sacrificing the contiguity. These configurations are located at the top left region of Figure 4. The default parameter set for Stash corresponds to the optimized configuration chosen from the Pareto frontier, as shown in Table 1. Supplementary Figure 2 displays the same parameter sweep results on the Shasta assembly. It is worth noting that many of the well-performing parameter configurations.

**Table 1.** The empirically chosen space for *StashCut* parameters. The underlined values indicate values that were never seen on the pareto front, and the bolded numbers show the chosen default *StashCut* parameter set (refer to Figure 4).

| Parameter | Explored Values |
| --- | --- |
| Number of Frames | 2, **4**, 6, 8, 10 |
| Stride (bp) | <u>1</u>, <u>20</u>, **40**, 60, <u>80</u> |
| Delta (bp) | <u>1</u>, <u>10</u>, **100**, 1000, 10000 |
| Threshold | 17, 18, 19, **20**, <u>21</u> |
| Kernel Radius (bp) | 1, **2**, 3, 4, <u>5</u> |

**Fig. 4.**
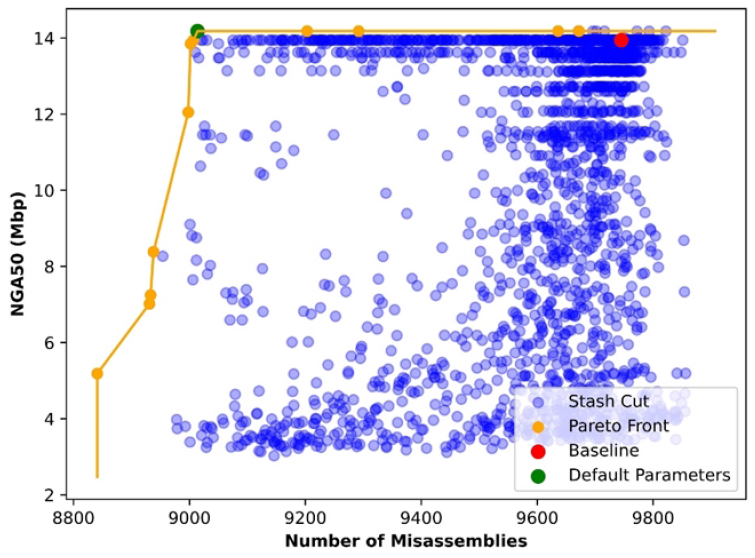
*StashCut* applied with 3,125 different configurations on the Flye assembly. Results were obtained using QUAST after scaffolding the assemblies using ntLink. As displayed by the Pareto frontier, higher quality cuts result in lower number of misassemblies and higher NGA50 length.

To understand the effect of each Stash parameter, we color-coded the data points from Figure 4 and show them in Figure 5. In this figure, a higher density of the configurations is on the top left region, and brighter points are shown on top of the darker points. As suggested in Figure 5.A, the *number of frames* parameter should be kept under five to achieve the highest performance. This is sensible as a high *number of frames* will cause the window maximum operator to fade out the details of the matches signal. Moreover, Figure 5.B indicates that low *stride* values (<~20) are not effective, suggesting that a window that spans over a larger genomic region might be more resistant to sequencing errors. For example, if a frame of a window contains an indel error, with a low *stride*, the following frames of the same window will also include that error, and therefore, resulting in a Stash cut in the region, reducing the contiguity without cutting at a true misassembly. Figure 5.C suggests higher cut thresholds, which would result in higher contiguity as a result of lower number of false positive cuts. It is worth mentioning that the *threshold* parameter is dependent on the *number of frames* parameters. A good *threshold* value is defined based on the expected number of matches between a pair of unrelated windows. Figure 5.D combines *threshold* and *number of frames* to show this dependency, and indicates that high thresholds paired with low *number of frames* and low thresholds paired with high *number of frames* will not result in the optimum cut. Supplementary Figures 3 and 4 further demonstrate the optimal *threshold* value through precision-recall curves which are generated using custom definitions for precision and recall (Supplementary Method 1). Figure 5.E does not show an obvious pattern for the *delta* parameter. The only consideration for this parameter is that the value needs to be set with regards to the minimum contig length. As shown in Figure 5.F, higher *kernel* radius values result in the same generalization and loss of details as high *number of frames* values. As can be seen from Figure 6 and detailed in Supplementary Table 5, *StashCut* with the default parameter set (Table 1) is comparable to both Tigmint-long and Inspector. For the Flye assembly of human cell line NA24385 data, *StashCut* detects 13% of the extensive misassemblies without decreasing the NGA50 length metric. On the same dataset, Inspector results in insubstantial changes to both NGA50 and the number of misassemblies (< 0.3%), and Tigmint-long results in 2% reduction in misassemblies with the downside of reducing the NGA50 by 22%. By performing ntLink scaffolding on the baseline Flye assembly and the output of each misassembly correction module, we can see that the scaffolded *StashCut* assembly still has 7.6% fewer the misassemblies compared to the baseline scaffolded Flye (9,003 vs. 9,745), which is considerably higher than the Inspector (−0.1%) and Tigmint-long (2.6%) runs. However, this improvement comes at the cost of lower NGA50 compared to the other methods (StashCut: 14,181,592 bp, Inspector: 14,624,442 bp, Tigmint-long: 16,500,644 bp) as shown in Supplementary Table 6. When correcting the Shasta assembly, *StashCut* corrects more extensive misassemblies than Inspector (by 239) and Tigmint-long (by 337) and produces a superior scaffolded assembly considering the NGA50 metric (*StashCut*: Similar to Flye, Supplementary Tables 8 and 9 provide details of the runs for Shasta.

**Fig. 5.**
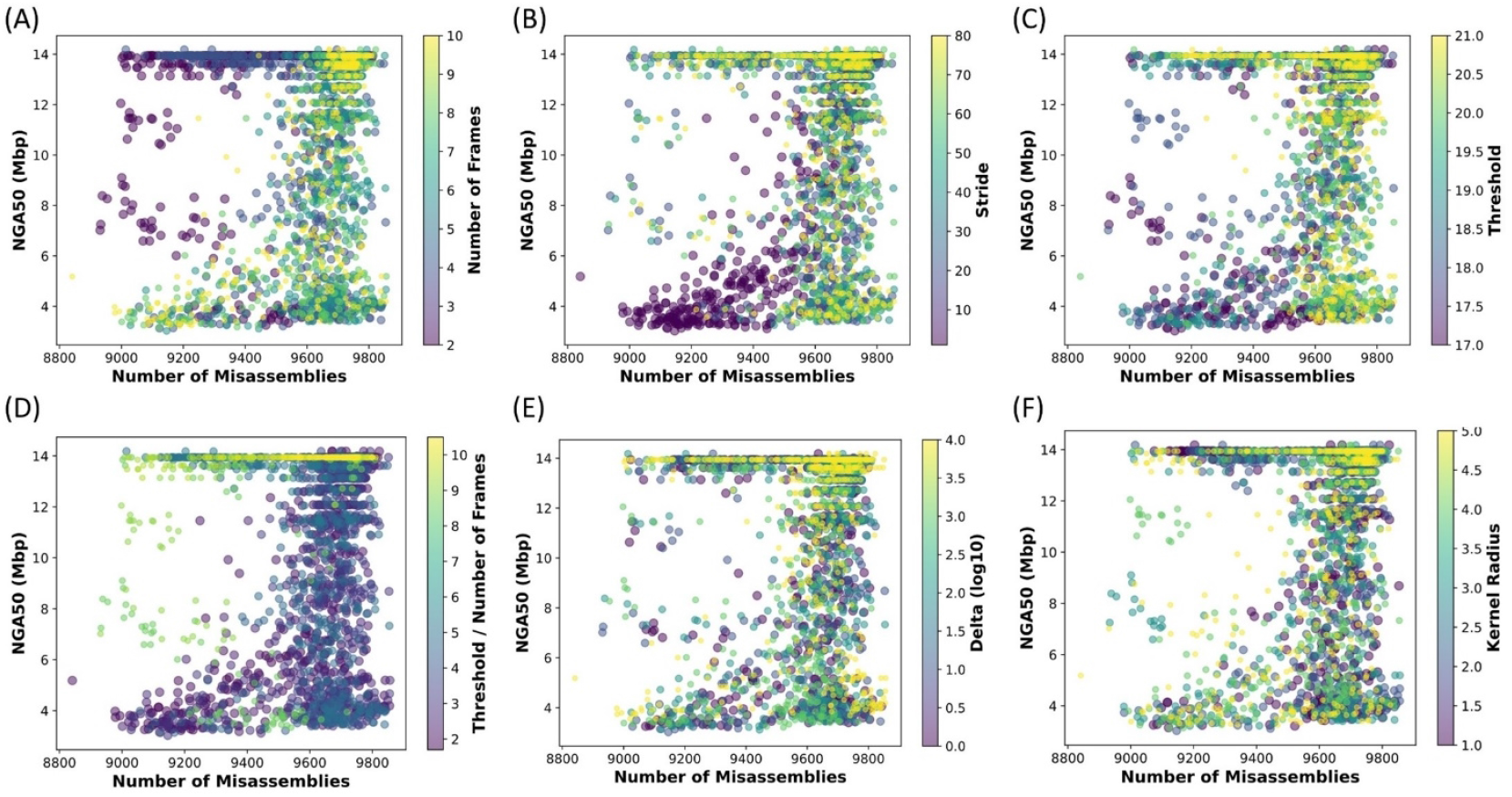
The effect of each Stash parameter on the scaffolded result after *StashCut*. Some parameters are easy to justify, while others do not seem to have an immediately obvious pattern. Panel (D) represent both the *threshold* and the *number of frames* parameter. For improved visibility, points with higher color values are shown in front while points with lower color values have larger sizes.

**Fig. 6.**
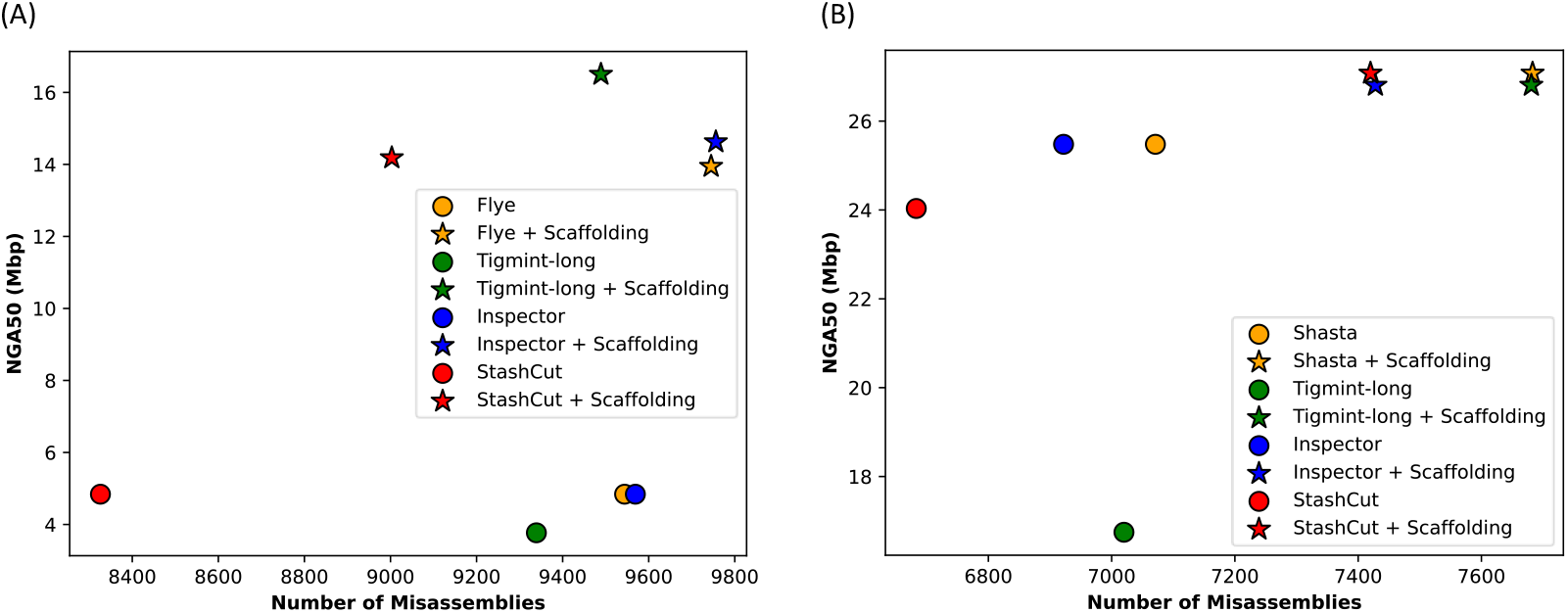
Performance comparison between *StashCut* and other misassembly correction methods on Flye (A) and Shasta (B) ssemblies. Each figure contains two set of evaluations; circles represent the corrected assembly while stars include an dditional scaffolding step after correction. *StashCut*, represented in red, achieves the least number of misassemblies both efore and after scaffolding, and Tigmint-long, represented in green, achieves the most contiguous scaffolded Flye ssembly. On the Shasta assembly, *StashCut* results in an objectively better scaffolded assembly than the baseline shown n orange.

To complement our results, we also evaluated StashCut’s performance using Oxford Nanopore Q20+ reads from the publicly available GM24385 (HG002) dataset, released by Oxford Nanopore Technologies. While ONT reads generally exhibit higher error rates—especially indels in homopolymer regions—they offer ultra-long read lengths and continue to gain popularity in *de novo* assembly workflows. The ONT-based results, including both contig-level and scaffold-level evaluations, are presented in the Supplementary Results 1.

In addition to human genome datasets, we further evaluated StashCut’s performance on a non-human model organism. Using ~428-fold coverage PacBio RSII reads from *C. elegans* (SRR7594465), we applied StashCut to an existing assembly (ASM1813679v1) and assessed its performance using the WBcel235 reference. This experiment demonstrates StashCut’s applicability beyond human genomes and across different long-read sequencing platforms. A detailed summary of these results is provided in the Supplementary Results 2.

Filling the Stash with the sequencing data used in our experiments takes 292 minutes using 8 threads and consumes a peak memory of 8 GB. Table 2 shows the benchmarking results for all three misassembly correction modules on the Flye assembly using 8 threads. The default configuration of StashCut, with number of frames equal to 4, runs 6.5-fold slower but uses about half the memory compared to Tigmint-long. It runs three times faster and uses two-thirds less memory compared to Inspector. The table also includes the benchmarking results for a *StashCut* with number of frames equal to 1 in order to show that the time complexity of *StashCut* is quadratically proportional to the *number of frames* parameter.

**Table 2.**
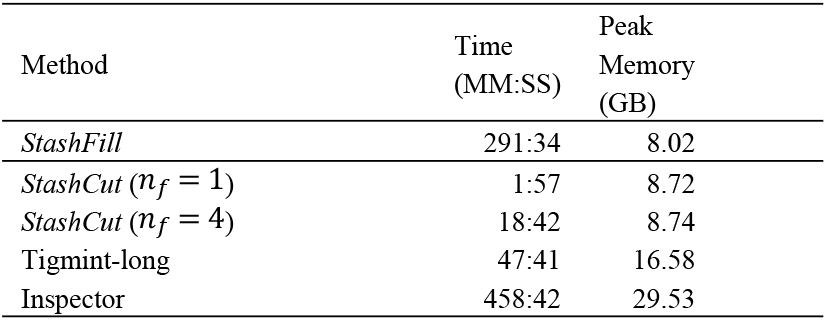
Benchmarking results for Stash, Inspector, and Tigmint-long applied on the Flye assembly using 8 threads. Note that the complete Stash pipeline requires both *StashFill* and *StashCut*.

| Method | Time<br>(MM:SS) | Peak<br>Memory<br>(GB) |
| --- | --- | --- |
| *StashFill* | 291:34 | 8.02 |
| *StashCut* ($n_f = 1$) | 1:57 | 8.72 |
| *StashCut* ($n_f = 4$) | 18:42 | 8.74 |
| Tigmint-long | 47:41 | 16.58 |
| Inspector | 458:42 | 29.53 |

## 4. Discussion

The ability to implicitly store an arbitrary property (such as coverage) from a large set of inputs (like a sequencing read set) is a primary contribution of Stash, making it a useful tool for bioinformatics applications, especially for the (B) genome assembly pipeline. We show how a StashCut-corrected assembly leads to a higher-quality post-scaffolding result due to reduced number of misassemblies. Compared to the other state-of-the-art misassembly detection methods such as Tigmint-long or Inspector, Stash does not rely on any sequence alignment; moreover, it has a linear time complexity with respect to the input data size and a bounded memory usage set by its user-defined parameters, making it computationally efficient.

Stash presents another notable advantage which is not being confined to a specific input read length category. We anticipate its efficacy across short reads, linked reads, and long reads. For short reads, the same filling and querying approach can be used, and for linked reads, the associated read barcodes can be hashed and stored instead of read IDs. Our findings in this study demonstrate the robust performance of Stash within long-read assembly pipelines due to their long-range information. This is important as the continuing progress in accuracy, throughput, and cost reduction have made long-read sequencing useful in genomics.

The quality of the reads has a direct effect on the performance of Stash, as the extracted number of matches signal can be vulnerable to sequencing errors. This is because of the low coverage of overlapping reads represented by low number of matches in regions with indels, gaps, and mismatches. In our experiments, we primarily used ~30-fold coverage PacBio HiFi long-read sequencing data from the human cell line NA24385, with an average Phred quality score of 35, consistent with high-accuracy long-read sequencing (Ewing, et al., 1998) (Ewing & Green, 1998). PacBio HiFi reads are particularly well suited for misassembly detection due to their low error rates. Additionally, the uniform coverage of these input reads is sufficient to saturate Stash frames, considering the allocated frame dimensions in memory. Due to this sensitivity to sequencing errors, we expect the number of matches signal extracted from Stash to have lower number of false positive drops when the data structure has been filled with high-quality short reads.

In the conducted experiments, Stash reduces the number of misassemblies in an input assembly of interest without noticeably decreasing the contiguity measured by the NGA50 or NG50 metrics. In our benchmarks, we filled the data structure with the input sequencing reads and applied the *StashCut* module to the corresponding long read assemblies generated by the Flye and Shasta assemblers. By breaking misassemblies, the resulting subsequences can potentially be correctly joined during subsequent scaffolding. We passed the post-*StashCut* assembled genome to ntLink scaffolding and used the output to assess the quality of *StashCut* in the *de novo* assembly pipeline.

Regarding future work, there are several avenues that could be explored. First, further signal processing techniques can be applied on the number of matches signal in order to help identify genome misassembly positions. Moreover, machine learning approaches for signal processing can be performed on Stash fingerprints, to uncover patterns associated with misassemblies and improve detection accuracy. Also, as Stash acts as a probabilistic model, by using ensemble-based approaches such as combining multiple Stash instances, one can possibly increase the accuracy of the method even more depending on the application of interest. Finally, it is worth delving into other applications of Stash, including its potential in genomics research domains such as scaffolding, phasing, or read binning for metagenome assembly.

To sum up, Stash is a novel hash-based data structure used for storing and retrieving large amounts of sequence mapping information such as genome sequencing reads using a lossy representation, allowing for the implicit storage of a specific property associated with the input observations. The successful utility of Stash in the misassembly detection problem has been explored when the stored property is *coverage by a read set*.

## Acknowledgements and Funding

The authors would like to acknowledge discussions with Amirhossein Afshinfard on statistical properties of Stash and thank René Warren for providing feedback at different stages of the project.

This study is supported by the National Institutes of Health [2R01HG007182-04A1] and Canadian Institutes of Health Research (CIHR) [PJT-183608]. The content of this article is exclusively the responsibility of the authors and does not represent the official views of the National Institutes of Health or other funding organizations. The funding organizations did not have a role in the study design, the collection, analysis, and interpretation of the data, or in writing the final manuscript.

## Data Availability

The accession codes of the sequencing datasets used for assembling the draft genomes or showcasing Stash algorithm are listed in Supplementary Table 1.

